# Complex effects of chytrid parasites on the growth of the cyanobacterium *Planktothrix rubescens* across interacting temperature and light gradients

**DOI:** 10.1101/2022.02.24.481659

**Authors:** Joren Wierenga, Mridul K. Thomas, Ravi Ranjan, Bas W. Ibelings

## Abstract

Chytrids are important drivers of aquatic ecosystems as phytoplankton parasites. The interaction between these parasites and their hosts are shaped by abiotic factors such as temperature and light. Here, we performed a full-factorial experiment to study how temperature and light interact to affect the dynamics of the bloom-forming toxic cyanobacterium *Planktothrix rubescens* and its chytrid parasite. We used a dynamic host-parasite model to explore how temperature and light affect long term dynamics. At low temperatures, chytrids do not survive. Higher light and temperature levels stimulated both phytoplankton and chytrid growth, with complex effects on their dynamics. Model exploration indicates that increasing temperature and light shifts equilibrium outcomes from *P. rubescens* persisting alone to stable coexistence and then to limit cycles. This provides an alternative biological explanation for why *P. rubescens* is mainly found in the relatively cold and dark lake metalimnion – it may enable avoidance of its parasite. Our study emphasizes the importance of investigating how abiotic factors interact with biotic interactions to drive complex outcomes.

## Introduction

Cyanobacteria are important primary producers in aquatic systems [1] that can be harmful when they form dense blooms, which can sometimes contain high concentrations of toxins [2-4]. *Planktothrix rubescens* is a common cyanobacterium that often produces toxic blooms, with high concentrations of hepatotoxins like microcystins or neurotoxins like anatoxin-a [5, 6]. It is typically found in the metalimnion of deep stratifying lakes, and performs better at lower light and temperature conditions than other cyanobacteria [7 - 11]. It has an efficient light-harvesting complex composed of phycocyanin and phycoerythrin in addition to chlorophyll, which allows it to photosynthesize at these low light levels [12, 13]. But even though *P. rubescens* can grow at low light and temperature, its growth rate increases with temperature and light, and is highest at values well outside the range they typically experience in their natural habitats. Why *P. rubescens* is rarely seen in high-light, high-temperature conditions that would appear to favour its growth remains an open question. One possible explanations that we investigate here is that temperature and light shape biotic interactions such as parasitism.

Phytoplankton growth rates increase with light and temperature up to optimal values [14], and decrease thereafter – slowly for light, very rapidly for temperature i.e. they are oppositely skewed unimodal functions [15-17]. The optimal temperature and light conditions for *P. rubescens* are poorly constrained based on existing experiments, but Oberhaus et al. (2007) measured growth rates at 15 & 25 °C and showed that it was faster at 25 °C. This implies an optimum temperature close to or above 25 °C because growth rate decreases rapidly above the optimum temperature [17]. Similarly, Bright & Walsby (2000) and Oberhaus et al. (2007) found no meaningful decrease in growth till the maximum irradiance levels they measured in white light (200 and 300 μE m^-2^ s^-1^) [13, 18]. The optimum may be lower than this at 10-15 °C, but the measured difference in growth rate was well within the range of experimental error and an optimum above 200 μE m-^2^ s^-1^ cannot be ruled out at these temperatures either. But in lakes, the species appears to be most abundant at temperatures of 10-15 °C, although there is based on limited data; abundance peaks from 6.5 to 20 °C have been seen [7, 19-22]. And it is often found at the depth where irradiance is 0.1% – 1% of that at the surface (∼ 1 – 20 μE m^*-2*^ s^*-1*^). They maintain themselves at this depth through light-mediated buoyancy regulation, with lift provided by gas vesicles offset by carbohydrate ballast [23].

The dynamics of cyanobacteria are driven not just by abiotic factors such as temperature and light, but also by biotic interactions with predators and parasites [24-26]. Parasites are important but neglected biotic drivers of ecological dynamics, commonly affecting cyanobacteria as well as other species of phytoplankton [27-30]. Parasites have been shown to impact host populations of primary producers, with knock-on-effects at higher trophic levels and community structure [31-35]. One of the most ubiquitous groups of aquatic parasites is the Chytridiomycota, a large and diverse group of fungi that are best known for the role they are playing in amphibian extinctions worldwide [36-38]. Chytrids have complex life cycles (described in [39]) and also infect many species of algae and cyanobacteria, including *P. rubescens*. They are often involved in the termination of blooms of cyanobacteria and other phytoplankton [40-44]. Chytrids can act as a shunt in aquatic ecosystems, transferring nutrients and carbon to higher trophic levels through infection of phytoplankton that are otherwise resistant to grazing [45-47].

The chytrid parasite is also affected by temperature and light, not just through the direct effects of temperature on its physiological processes, but also through temperature and light effects on the host [9-11, 48, 49]. For example, light affects the release of dissolved organic carbon (DOC) by phytoplankton [50, 51], which subsequently impacts chemotaxis of chytrid zoospores, and thus infection dynamics [9, 11, 52, 53]. However, we know little about how light and temperature interact to affect chytrid infections generally, and even less in the case of *P. rubescens*. Much of the work on chytrid – host interactions has been done on the diatom *Asterionella formosa* and its associated chytrids, *Rhizophydium planktonicum* and *Zygorhizidium planktonicum*. In these host-parasite pairs, both low temperature and low light can (independently) provide refuge for the host [9, 52]; ‘refuge’ in this case indicates a set of environmental conditions that prevents chytrid infection and not a physical location. *Asterionella* appears to have both a low-temperature and a high-temperature refuge, indicating that the physiological tolerance range of chytrids is narrow relative to at least some phytoplankton [48]. For *P. rubescens*, only low temperatures have been shown to offer a refuge from chytrid infections with high temperature not investigated [10]. The closely-related species *Planktothrix agardhii* does appear to have a high-temperature refuge; McKindles et al. (2021a) found that chytrid infection rates (measured as % increase in filaments infected per day) declined above approximately 22 °C to nearly zero by 29 °C [49]. This environmental sensitivity of the chytrids has important consequences for *P. rubescens*, which is believed to experience substantial variation in temperature and light due to its typical habitat in the metalimnion. Understanding how the environment shapes the dynamics and location of *P. rubescens* requires us to examine how these abiotic factors interact to shape its growth and biotic interactions. The size and complexity of such experiments means that they are rarely performed, weakening our ability to understand how environmental change affects natural populations and communities.

Here we investigated how light and temperature interact to shape *Planktothrix-*chytrid interactions using a combination of experiments and theory. We performed a full-factorial experiment that features four temperatures, four light levels and two infection statuses (infected and uninfected *P. rubescens* cultures). We extended a dynamic host-parasite model [54-57] to explore the consequences of temperature-light interactions for *P. rubescens*-chytrid dynamics. To examine this specific system, we made model parameters dependent on temperature and light and estimated them from our experiment where possible. We investigated whether the thermal refugium for *P. rubescens* depends on light intensity, indicating that abiotic interactions shape these ecologically important biotic interactions. We demonstrate that increasing light and temperature stimulated both phytoplankton and chytrid growth, with complex effects on host-parasite interactions.

## Materials and Methods

### Experimental design

We used a full factorial experimental design with 4 temperatures (6, 11, 16 and 21 °C), 4 light levels (2, 7, 14 and 21 μE m^-2^ s^-1^) and 2 infection status levels (infected/uninfected *P. rubescens* cultures), for a total of 32 treatment combinations. We used 4 replicates at each combination, for a total of 128 experimental units. Fig. S1 shows a schematic illustrating the experimental design.

### Culture details and experimental conditions

*Planktothrix rubescens* strain NIVA-CYA98 and the chytrid parasite Chy-Kol2008 (Rhizophydiales) that were used in this study are monoclonal but non-axenic; bacterial biomass was kept low by semi continuous growth, confirmed by microscopy. WC medium was used to grow *P. rubescens* [58], modified to exclude silicon as a macronutrient.

Before the start of the experiment the *P. rubescen*s cultures were acclimatised to the different experimental conditions. Temperature was regulated by using heated water baths in a refrigerated room at 4 °C, logged with calibrated temperature loggers (HOBO Onset, UX120-006M). Light was provided by fluorescent tubes (Osram neutral-white, 4000K), and light intensities were manipulated using neutral density filters shading the culture flasks (manufactured by Lee Filters, Hampshire UK, filters numbers 211, 210 and 209). Light intensities were measured inside the shaded flasks (Licor LI-250A light meter, US-SQS/WB quantum sensor). Photoperiod was set to a 16:8 hour cycle of light and dark. The experiment was performed in 250 mL culture flasks (Greiner Bio One, Item No.: 658195). Cultures were shaken daily during acclimatisation and during the experiment.

At the start of the experiment, the culture flasks were filled with 100 mL of *P. rubescens* suspension, diluted to a biovolume of 4 nL mL^-1^, so all treatments had the same density at the start of the experiment. Subsequently, half of these flasks inoculated with a dense zoospore suspension, constituting the infected treatment. The zoospore suspension was made by filtering a heavily infected culture (16 °C, 11 μE m^-2^ s^-1^) of *P. rubescens* through a glass serological pipette filled with loosely packed glass fibers that had been autoclaved. The glass fiber filter that is created this way lets through most of the zoospores but is highly efficient in removing the filaments of *P. rubescens*. One mL of this dense zoospore suspension was added to the infected treatment, resulting in 70 zoospores mL^-1^ at the start of the experiment. The experiment lasted for 20 days.

### Sampling and imaging

2 mL samples were taken every other day during the course of the experiment and fixed in 0.5% final concentration glutaraldehyde. The samples were stored in 2 mL sample tubes at 4°C.

To analyze the samples from 128 experimental units, a high throughput biovolume measurement method was developed (Wierenga et al., in prep). The method relies on high resolution composite images of the entire surface of a clear bottom well, imaged by an automated microscope. 300 μL of each sample was pipetted in duplicate into a clear bottom 96-well plate (Greiner Bio one Item No: 655096) for image analysis. The plates were put in the fridge for 24 hours, to allow enough time for settling of the filaments. Using a Biotek cytation-3, image composites were taken of the entire surface of each well bottom with a 4x magnification objective. Images were taken in the chlorophyll and phycoerythrin autofluorescence channel (586 nm excitation and 647 nm emission). This yields high quality images with bright filaments against a dark background, ideal for image analysis. For infection measurements 300 μL fixed sample was pipetted in duplicate into clear-bottom 96-well plates. The plates were put in the fridge for 24 hours, to allow enough time for settling of the filaments. To visualize the chytrid sporangia, 10 μL calcofluor-white (CFW) was added to each well to get a 5 μg/mL final solution. To increase fluorescence output of the CFW stain, 10 μL of sodium hydroxide (pH > 13.5) was added to increase the pH in the well. After incubation for 10 minutes, the plates were imaged with the Biotek cytation-3, in three different channels: brightfield, chlorophyll and phycoerythrin autofluorescence (586 nm excitation, 647 nm emission), and CFW (377 nm excitation, 447 nm emission). These composite images were combined to generate images which clearly show the outline of the filaments and stained sporangia. Furthermore, the viability of filaments can be assessed from the autofluorescence signal in the images.

### Image analysis

To measure the biovolume, the autofluorescence image composites were analyzed with the open-source image analysis tool *ImageJ*, using the plugin *skeletonize*. In short, this method uses a “Skeletonization algorithm” to compute the total length of all filaments in a picture. These are subsequently converted to biovolume using the equation for calculating the volume of a cylinder (*V* = *πr*^2^*l*), where *r* is the mean radius of filaments and *l* is the total length of all filaments. The method is robust and can handle relatively high concentrations in samples because overlapping filaments is not a big problem. It requires only limited human input, reducing bias. Furthermore, with this method a relatively big sample of 300 μL is measured in its entirety, compared to manual counting of only a fraction of that volume, thereby improving precision.

To measure infections during the experiment, in the composite infection images a minimum of 50 filaments were measured and marked as infected or uninfected. Instead of using the frequency of infected and uninfected filaments to calculate the prevalence of infection, we used the length of infected and uninfected filaments. This results in a better representation of the prevalence of infection because there is a large variation in length, and moreover, infected filaments are typically shorter than uninfected filaments (as also seen in [59]).

### Data analysis

Data analysis and graphing was done using R version 4.0.4 and RStudio [60, 61]. ggplot2 (version 3.3.5) was used to make the graphs and the data processing was done with the help of the tidyverse packages (version 1.3.1) [60-63]. Comparisons of means were done using the robust yuen t-test from the package PairedData version 1.1.1 [64].

### Dynamical model

We used a dynamical host – parasite model to assess the dependence of chytrid – phytoplankton dynamics on temperature and light. We modified a model from Frenken et al. (2020) [55] by incorporating temperature-and light-dependence of the parameters. The model consists of three equations, which track the dynamics of uninfected hosts (*H*_*u*_), infected hosts (*H*_*i*_) and free-swimming zoospores (*Z*).

The uninfected host *H*_*u*_ follows logistic growth with a carrying capacity *K* (nL ml^-1^) and a growth rate *r* (day^-1^). Losses of uninfected host are defined by a density-independent mortality rate: *m*_*u*_ (day^-1^), and by infection of uninfected host by a zoospore, which converts uninfected host into infected host. The infectivity constant *I* (mL cell^-1^) is a measure for the infection efficiency of zoospores (mL cell^-1^).

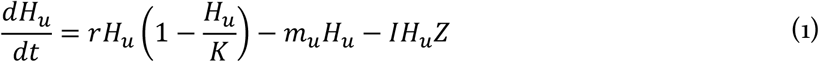

Infected hosts *H*_*i*_ increase with every uninfected host that gets infected by the attachment of a zoospore. This is the same element as the loss due to infection for the uninfected host (*IH*_*u*_*Z*). Infected host decreases with a density-independent mortality rate *m*_*i*_ (day ^-1^) and with maturation of sporangia given by the development time *τ* (days). The maturation of sporangia assumes infected host *H*_*i*_ is converted into zoospores.

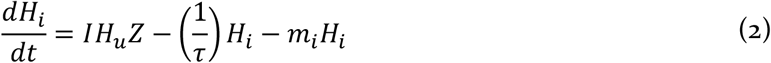

Chytrid zoospores *Z* increase when sporangia develop with development time *τ* and release new zoospores. The zoospore production parameter *ρ* (cells nL^-1^) defines the number of spores produced per biovolume of infected host. Spores are lost when they infect an uninfected host, and additionally at a density-independent mortality rate *m*_*z*_ (day ^-1^).

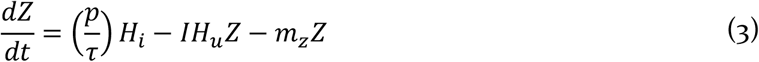

*Model parameter values*: Some model parameters (Table 1) were calculated directly from our experimental data: the infectivity parameter *I* and the growth rate *r*, making them functions of light and temperature. The infectivity parameter *I* is calculated by assuming exponential decay of *H*_*u*_ with *Z*, and is directly calculated from experimental data according to the following equation:

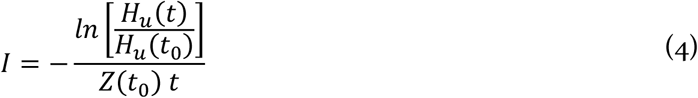

where *t*_*0*_ is the beginning of the experiment and *t* is the length of the chosen period over which *I* was calculated (2 days; see supplementary information for details). The phytoplankton growth rate *r* was determined by fitting an exponential model to *P. rubescens* density over time in the uninfected treatments. To run the dynamical model at a higher resolution for temperature and light than the intervals used for the experiment, a generalized additive model (GAM) was fitted to calculated values of *r* and *I* across the temperature and light levels used in the experiment. This GAM-model was subsequently used in the dynamical host – parasite model to generate values for *r* and *I* at interpolated temperature and light. The parameters *K* and *ρ* are kept constant. The carrying capacity is set at 600 nL mL^-1^, based on a logistic fit through experimental data at high light and temperature. The production of zoospores is *ρ* calculated from supplementary data from Frenken et al. (2020) [55], which has zoospore production data on a very closely related strain of *P. rubescens* and chytrids. It is calculated by dividing newly produced zoospores by the biovolume of the host 2 days prior, and is found to be 20 cells nL^-1^, which is set for each temperature and light combination. Infective lifetime of zoospores was found to be 2.71 days at 17 °C. This translates into a mortality rate *m*_*z*_ of 0.37 (day^-1^) which we set to exponentially increase with temperature. Development time for sporangia is found to be between 2 and 3 days based on routine observations of infected cultures and is set to exponentially decrease with temperature. The dynamical model was analyzed in R using the R Package: deSolve, version 1.28 [65].

**Table 1:** Overview of all parameters and variables used in the dynamical model. Where possible parameters were based on data from our experiment or from previous experiments with closely related organisms. The dependence of the parameters on temperature, light or both is indicated.

| Symbol | Shape/Dependency | Meaning | Value/equation | Units |
| --- | --- | --- | --- | --- |
| <u>Variables</u> |  |  |  |  |
| $H_u$ | | Uninfected host | | nL mL<sup>-1</sup> |
| $H_i$ | | Infected host | | nL mL<sup>-1</sup> |
| $Z$ | | Free-swimming zoospores | | cells mL<sup>-1</sup> |
| <u>Parameters</u> |  |  |  |  |
| $K$ | Constant | Carrying capacity | 600 | nL mL<sup>-1</sup> |
| $I$ | GAM with temperature and light | Infectivity | GAM model | mL cell<sup>-1</sup> day<sup>-1</sup> |
| $r$ | GAM with temperature and light | Growth rate of the uninfected host | GAM model | day<sup>-1</sup> |
| $m_u$ | Exponential increase with temperature | Death rate of uninfected host | $m_u = 4.62 \cdot 10^{-4} \cdot e^{0.0783 \cdot T}$ | day<sup>-1</sup> |
| $m_i$ | Exponential increase with temperature | Death rate of infected host | $m_i = 0.0305 \cdot e^{0.117 \cdot T}$ | day<sup>-1</sup> |
| $p$ | Constant | Zoospore production per biovolume of infected host | 20 | cells nL<sup>-1</sup> |
| $m_z$ | Exponential increase with temperature | Death rate zoospores | $m_z = 0.035 \cdot e^{0.128 \cdot T}$ | day<sup>-1</sup> |
| $\tau$ | Exponential decrease with temperature | Development time of zoospores in sporangia | $\tau = 16.42 \cdot e^{-0.099 \cdot T}$ | Time in days |

There is no straightforward way to calculate the mortality rates *m*_*u*_ and *m*_*i*_ from available data; we therefore assumed *m*_*u*_ to be equal to 1% of the maximum growth rate at each temperature (i.e. growth rate at the highest light level, 21 μE m^-1^ s^-1^). The mortality rate of infected *P. rubescens* is based on the same fit, with a 50 percent higher dependency on temperature and a higher initial value to account for the increased mortality due to infection. For more details about this temperature and light dependence, please refer to the supplementary information.

## Results

### Temperature and light effects on uninfected and infected Planktothrix growth rates

Temperature and light interacted to shape the growth of *P. rubescens* and its susceptibility to chytrid infections (Fig. 1, Table 1). The expected cold refuge [10] occurred in our experiments at temperatures of 11 °C and lower; no infections occurred at 11 °C and below. The growth rate of *P. rubescens* generally increased with both temperature and light at the levels tested in this experiment, but the shape of the interaction was complex. At low temperature, increasing light had little effect on growth. At low light, increasing temperature above 11 °C decreased growth rate (Fig. 1). Furthermore, the positive effect of temperature on the growth rate increased with light, whereas the positive effect of light on the growth rate, increased only till 14 μE m^-2^ s^-1^ (Fig. 1D, lines on top of each other).

**Figure 1.**
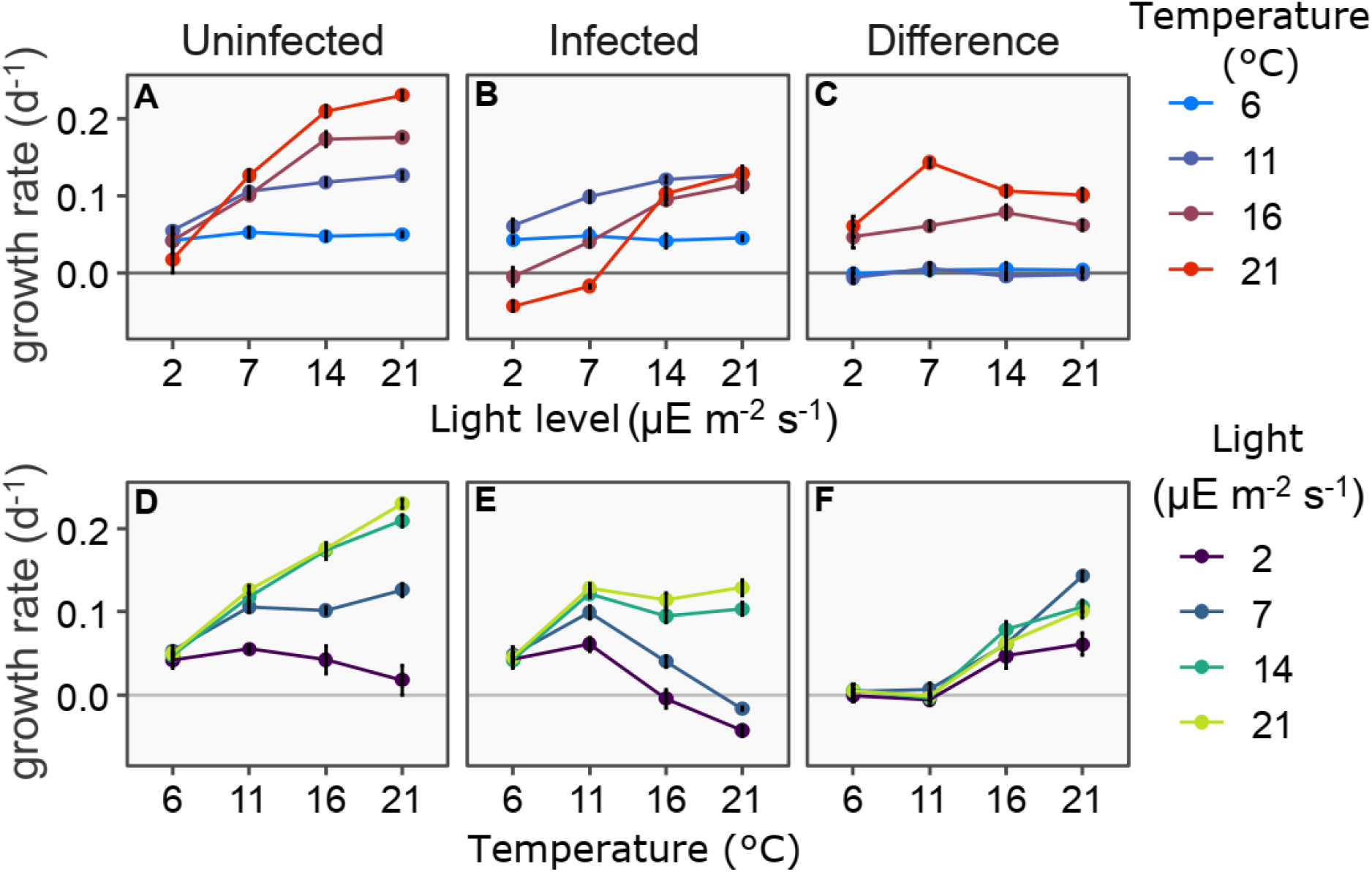
*Planktothrix* growth rate varies strongly with temperature, light and infection status (**A**) Growth rate of uninfected and infected *P. rubescens* as a function of light at each of the 4 temperatures tested. The effect of the interaction between light and temperature is clearly visible both in infected and uninfected treatments. Infection reduces growth rate at 16 and 21 °C but not at 6 and 11 °C. Within each temperature, changing light has little effect on the difference in growth rate between infected and uninfected treatments. (**B**) The same data, viewed with temperature on X-axis and with light levels indicated by color. Within each light level, changing temperature has a large effect on the difference in growth rate between infected and uninfected treatments. In all panels dots represent mean growth rates and error bars represent the standard error of the mean.

### Effects of chytrid infection on Planktothrix growth

Chytrids did not strongly affect the growth rate of *P. rubescens* at temperatures of 11 °C and lower. The growth rate remained essentially unchanged between the infected and uninfected treatment, confirming the refuge for *P. rubescens* at low temperatures (Fig. 1C). At temperatures of 16 °C and higher, the chytrid infection led to a substantially lower growth rate at all light levels tested. The reduction is similar across light levels (Fig. 1C). The biggest effect of the chytrid infection occurs at the highest temperature, with a big change in growth rate. This results in negative growth rates of *P. rubescens* at lower light levels (Fig. 1B, 1E). Growth of *P. rubescens* is very low at high temperature and low light (21 °C, 2 μE m^-2^ s^-1^); infection led to further deterioration, resulting in negative growth rates for *P. rubescens* in the infected treatment (21 °C, at 2 and 7 μE m^-2^ s^-1^ and 16 °C at 2 μE m^-2^ s^-1^).

### Temperature and light effects on chytrid growth and infection prevalence

Temperature and light interact to shape chytrid growth, as indicated by the varying prevalence of infection on the last day of the experiment (Fig. 2A), and the growth and development of the infected and uninfected fractions of the *Planktothrix* biovolume (Fig. 1, 2B). Temperature is a major factor for chytrid growth showing increased prevalence of infection, especially at high light (Fig. 2A), and increased growth of infected BV with higher temperature (Fig. 2B). The effect of temperature is stronger at light. At the lowest light level (2 μE m^-2^ s^-1^), the effect of temperature is absent, and the prevalence of infection is similar between 16 °C and 21 °C. Chytrid growth, as indicated by the growing fraction of infected *P. rubescens* biovolume, was only higher at 21 °C than at 16 °C with light levels at or above 14 μE m^-2^ s^-1^. Furthermore, growth of uninfected biovolume at 16 °C was higher than at 21 °C even though growth of *P. rubescens* in the uninfected treatment is higher at 21 °C than at 16 °C (Fig. 2B). This indicates that at 21 °C *P. rubescens* is more vulnerable to chytrid infection than at 16 °C, especially at higher light levels. At 21 °C increased light also leads to a greater increase of the prevalence of infection compared to 16 °C. We did not find a low light refuge for *P. rubescens*. On the contrary, our experimental results show a large reduction in the growth rate of *P. rubescens* at low light with chytrid infection, especially at high temperatures (Fig. 1).

**Figure 2.**
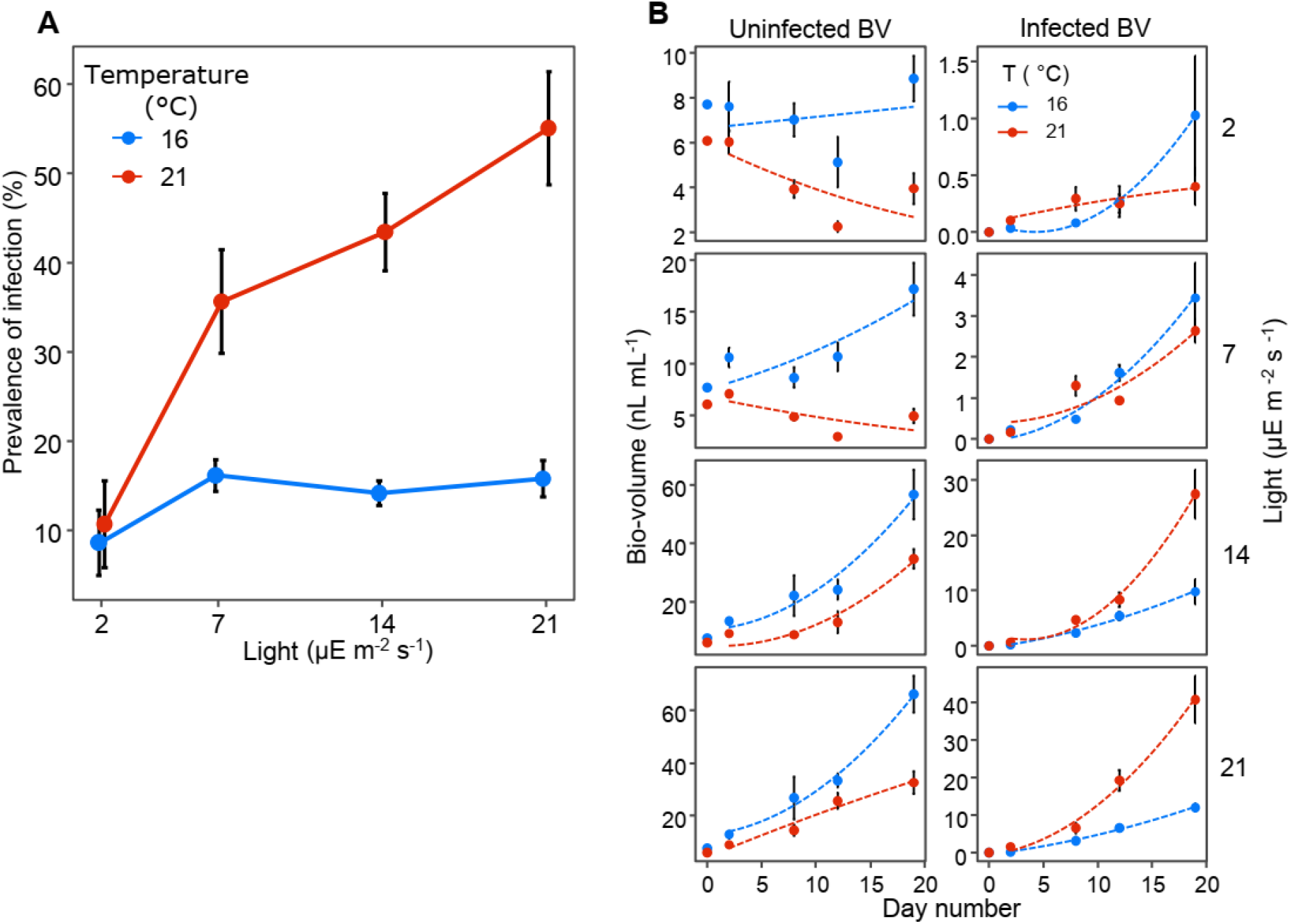
*P. rubescens* is affected more strongly by the chytrid parasite at 21 °C, especially at high light. (A) Prevalence of infection as a function of light level on the last day of the experiment. The prevalence of infection increases with increasing light at 21 °C, but not at 16 °C. At 6 and 11 °C no infected filaments were observed, and lines are not shown. (B) *P. rubescens* biovolume in the infected treatment, divided into uninfected and infected fractions; uninfected biovolume is shown in the left-and infected in the right-hand panels respectively. Asterisks indicate a statistically significant difference between means: * p<0.05, ** p<0.01

### Dynamical model

We investigated how temperature and light would interact to drive *P. rubescens* - chytrid dynamics using a host – parasite model. We see a clear refugium defined largely by conditions of low temperature (Fig. 3A), identified by the absence of zoospores (Fig. 3B) and the distinct outcomes of the model (the ‘only host’ region in Fig. 3C). The low temperature refuge is explained by the absence of chytrid infections at low temperature. This refuge is also dependent on light level; between approximately 13 and 15 °C, a refuge exists at very low light levels and at high (for this experiment) light levels, while intermediate light levels allow for persistence of the parasite. The highest densities of *P. rubescens* are found in refuge conditions and decrease with increasing temperature and light (Fig. 3A). The chytrid abundance is at its highest at an intermediate temperature between 14 and 16 °C, and light levels above 10 μE m^-2^ s^-1^.

**Figure 3.**
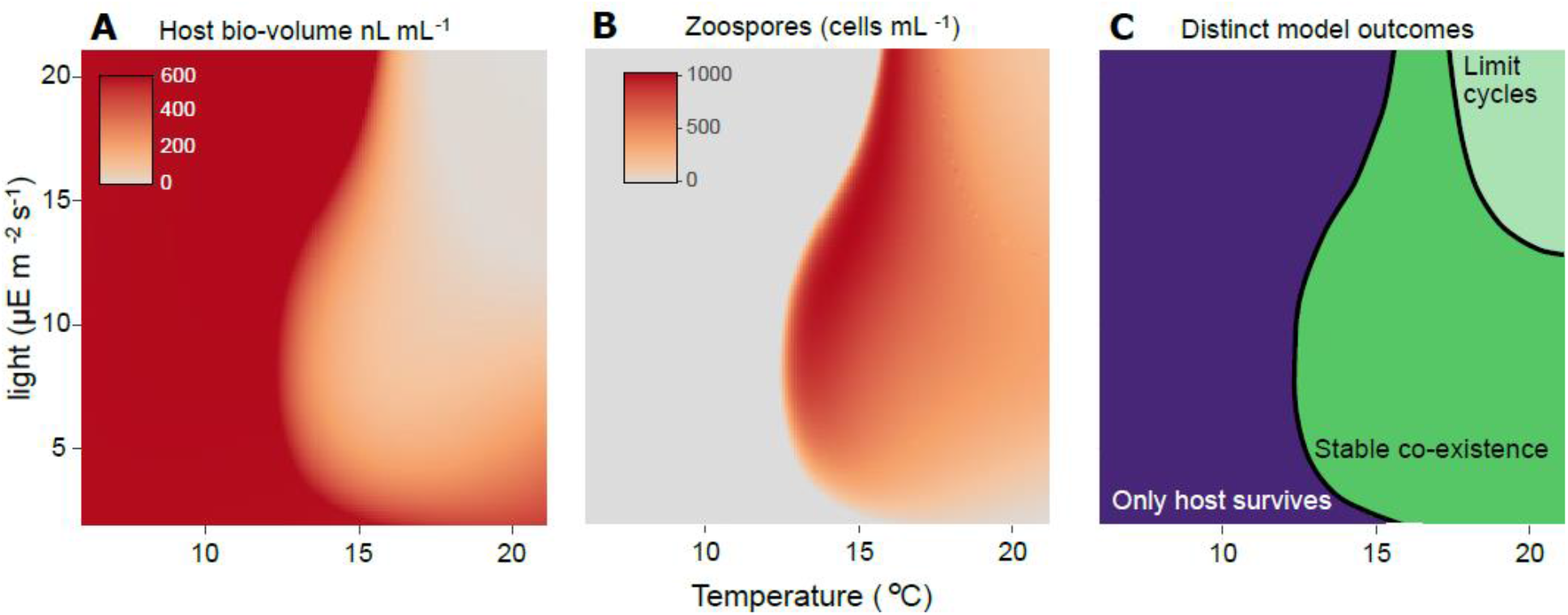
Outcomes of modelled host-parasite dynamics in constant temperature and light conditions from our theoretical model. Fig. 4 shows example time series from individual points on the temperature-light surface. (**A**) Host biovolume is greatest at temperatures below 13 °C and decreases above it. At high temperatures, host biovolume decreases with increasing light. (**B**) Chytrid zoospore abundance is highest at intermediate temperature and medium-to-high light levels. (**C**) The model shows 3 possible outcomes. The host persists alone at low temperatures, it exhibits limit cycles at high temperature and high light, and it coexists stably at intermediate conditions. Abundances shown in the limit cycle region are averages over the course of one cycle. Note that the model is intended to explore qualitative outcomes across parameter space and not to make precise quantitative predictions, for which additional factors (such as nutrients) would need to be considered.

Three distinct outcomes are observed at the different combinations of temperature and light we explored (Fig. 3C, Fig. 4). At low temperatures (< 13 °C), only the uninfected host can survive irrespective of light levels (Fig. 3A). The host (uninfected and infected) and the chytrid coexist stably at intermediate to high temperatures (Fig. 3C). However, the temperature range for stable coexistence is drastically reduced at high light levels (Fig. 3C). Finally, at high temperature and light levels, we find a limit cycle involving the uninfected host, infected host and the chytrid.

**Figure 4.**
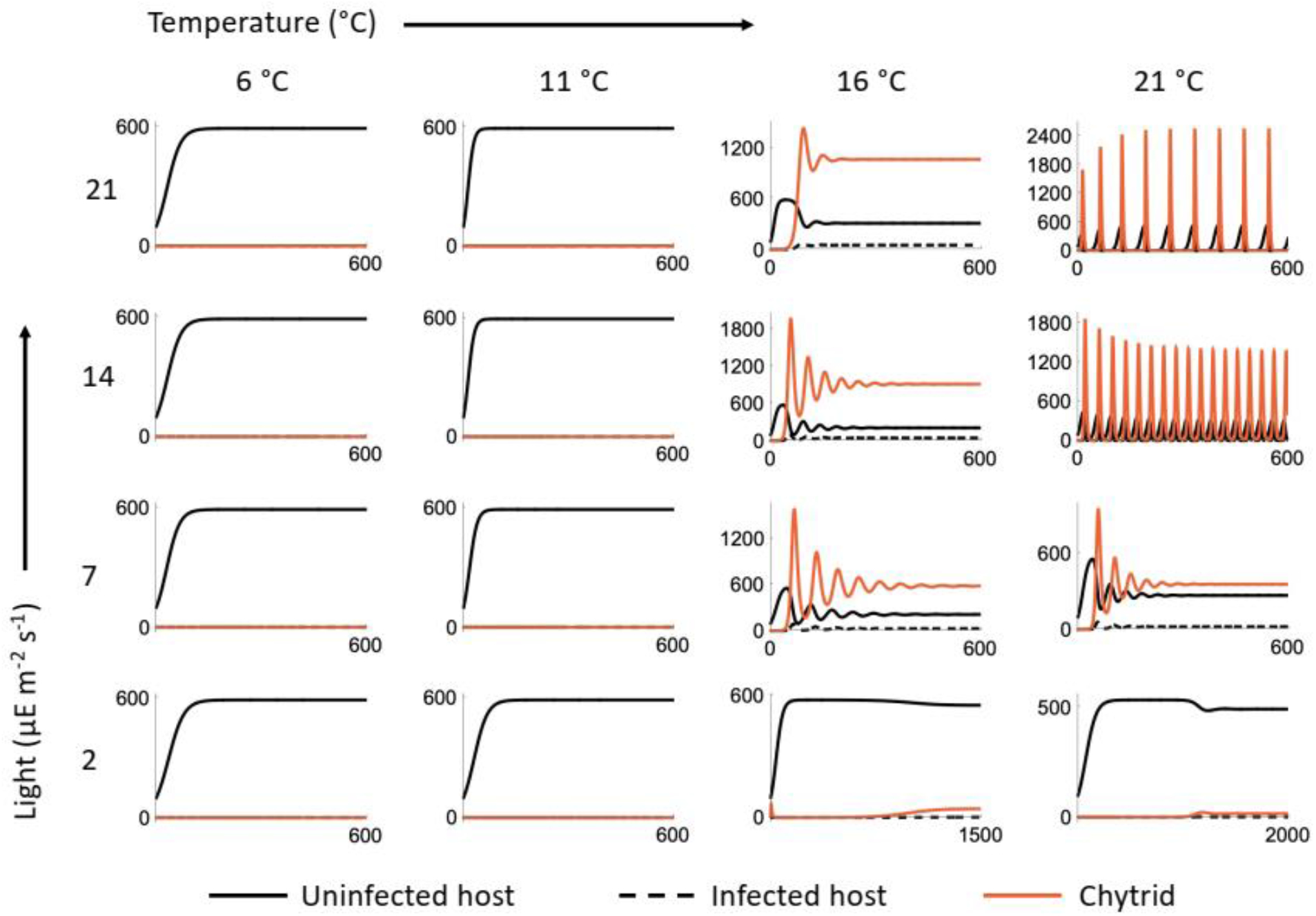
Example dynamics from the *P. rubescens* – chytrid model (summarized in Fig. 3), at the parameter combinations we measured experimentally. The solid and dashed black lines indicate uninfected and infected host densities respectively, while the red line indicates the chytrid density. At low temperatures, *P. rubescens* has a thermal refuge and its density reaches the carrying capacity. With increasing light and temperature there is a shift from stable coexistence (16 °C with 7, 14, 21 μE m^-2^ s^-1^, and 21 °C with 7 μE m^-2^ s^-1^) to limit cycles (21 °C with 14, 21 μE m^-2^ s^-1^).

## Discussion

### P. rubescens – chytrid growth and infection dynamics

We show that dynamics of *P. rubescens* and their chytrid parasites are heavily shaped by the interaction of temperature and light (Figs. 1-3). It has been previously shown that temperature and light are important factors [10, 11, 48, 49, 66, 67], but the effect of the interaction between temperature and light has not been well studied. To our knowledge, the only work on the interaction of temperature and light on chytrid-host dynamics is by Bruning [9], showing that in the diatom *Asterionella* and its chytrid parasite, that the two factors interact to affect infection dynamics. *P. rubescens* growth is most affected by chytrid infections at higher light and temperature levels. Even though we find relatively low infectivity of chytrids under low light, intrinsic slow growth due to low light means that chytrids can arrest *P. rubescens* growth. We thus do not find a physiological refuge for *P. rubescens* at low light as we hypothesized and was found for *Asterionella* [9].

### Indirect light resource utilization by chytrids

The chytrids in this experiment are obligate parasites [68], feeding on the host as a resource, and thus light is not a direct resource for the chytrids. Increased light does increase the release of dissolved organic carbon (DOC) by phytoplankton, affecting the affinity of zoospores to their host through chemotaxis and thus the interaction between the host and the parasite [9, 53, 69, 70]. However, here we find that at the lowest light level, increased temperature does not lead to increased prevalence of infection, and that increased light leads to a very small increase in the prevalence of infection at 16 °C. At 16 °C, temperature may be the limiting factor for chytrid growth, and more light does not lead to increased chytrid growth. At 21 °C however, the chytrid growth increases with light and thus might be resource-limited, with increased photosynthetic resources available at higher light intensity indicating indirect light utilization by acquiring energy resources from photosynthesis of the host while it is infected. Optimizing resource utilization by feeding while the host is actively photosynthesizing makes sense from an evolutionary perspective and is further suggested by the absence of a temperature effect at the lowest light level. This would not be expected if the chytrid uses only existing host resources at the time of infection for its reproduction and growth. Instead, it may feed off the host while alive, as occurs in most parasites [71]. A recent study has shown that increased light intensity and quality leads to bigger sporangia and increased chytrid transmission [11], indicating that chytrids indirectly utilize light through host photosynthesis. Another explanation could be that hosts grown under higher light stores more carbohydrates, and as such are a better energy source for the chytrid parasite.

### Dynamic model outcome and implications

The model shows three distinct outcomes over the range of conditions we explored: 1) only the host survives. 2) stable coexistence between the parasite and the host. 3) limit cycles of host and parasite. (Fig. 3C, Fig 4). Limit cycles occur at high temperatures and high light intensities (Fig. 3C, Fig. 4). Under these conditions, *P. rubescens* can grow rapidly but is vulnerable to infection, and infection under these conditions leads to a rapid decline of the host to very low densities. In a natural setting, chytrids are found both at low densities with only minimal impact on the host [72-74], as well as at high densities and high infection incidence that can rapidly terminate host blooms [34, 44, 66, 75]. In natural conditions, it is possible that periods of very low density could lead to local extinction of both host and parasite and so may not be congenial to the persistence of this host-parasite system despite the model outcome of limit cycles.

This suggests an interesting biotic alternative to abiotic factors explaining the niche of *P. rubescens*, which is almost always found around the 0.1 – 1% light level in stratified lakes [23]. The prevailing hypothesis is that they have adapted to a specific niche in the lake with relatively high nutrient levels, lower temperatures, and low light, where they outcompete other phytoplankton because they possess the accessory pigments phycoerythrin and phycocyanin [13]. However, this is inconsistent with the experimental data showing that growth rates of *P. rubescens* increase when temperature and light are increased beyond the limiting conditions found at the bottom of the euphotic zone; light levels up to 300 μE m^-2^ s^-1^, and temperatures up to at least 21 °C still promote growth [12, 13, 18, this study]. But at these high temperature and light levels, *P. rubescens* may be very sensitive to chytrid infections, and therefore unable to sustain growth, especially in competition with other phytoplankton species. Chytrid infection might thus limit *P. rubescens* to regions of the water column that are suboptimal for its growth, but safe from infection. If so, temperature, light and chytrids may together determine the vertical distribution of *P. rubescens* in stratified lakes. The model furthermore shows that the thermal refuge extends to higher temperatures at higher light levels, although we do not have experimental data to confirm this. We also cannot say whether a high-temperature refuge also exists as in the case of *P. agardhii* and *A. formosa* [48, 49]. Our explanation here for the *P. rubescens* vertical distribution would only hold if there was no such high-temperature refuge.

Although we believe this model offers us useful insights into the *P. rubescens* vertical distribution, its purpose is not to make precise quantitative predictions but instead to explore how temperature-light interactions may affect host-parasite dynamics. To make quantitative predictions, the model would at least need to account for several additional factors, including nutrient dynamics, temperature & light variation, and the effects of sinking, none of which we account for. Also, for some model parameter values, we used estimates based on the literature instead of measurements from our experiments (Table 1). These estimates are based on empirical data and physiological understanding, but there is unavoidable uncertainty about their true values. The values we chose may have led to lower prevalence of infection in the model (Fig. 3) than that seen in our experiments (Fig. 2), suggesting that some of these parameters could be improved upon if a predictive model is developed in future.

## Conclusion

We have shown that the host – parasite interaction of *P. rubescens* and chytrids is strongly affected by the interaction of temperature and light. Our results confirm the existence of a thermal refugium below at least 11 °C, and that this may vary with light intensity although there appears to be no specific low light refuge. In general, vulnerability of *P. rubescens* to chytrid infections increases with temperature and light. This may explain why *P. rubescens* is hardly ever found under warmer or high-light conditions, even though lab experiments show that these conditions lead to higher growth rates in the absence of chytrids. Our model does neglect some important complexities, such as the effects of nutrients on the biotic interaction, and the effects of light on host stoichiometry and consequently chytrid growth. Our results highlight how abiotic and biotic environmental factors can interact in complex ways to affect ecological dynamics.

## Acknowledgements

The authors gratefully acknowledge Thomas Rohrlack for early discussions which greatly helped putting us in the research direction described here, and Thijs Frenken & Takeshi Miki for discussions of their dynamic model and calculation of model parameters.

## Competing Interests

The authors declare no competing financial interests

## Data availability statement

The datasets generated during and/or analysed during the current study will be made available in the Zenodo repository on acceptance of this manuscript.

## Notes

### Competing Interest Statement

The authors have declared no competing interest.

### Summary of Updates

1) Introduction updated with more context. 2) Experimental design now shown in supplementary schematic figure. 3) Text updated for clarity. 4) A few details added to the methods section.

