## supplementary information for "Complex effects of chytrid parasites on the growth of the cyanobacterium *Planktothrix rubescens* across interacting temperature and light gradients"

### *I. Experimental design:*

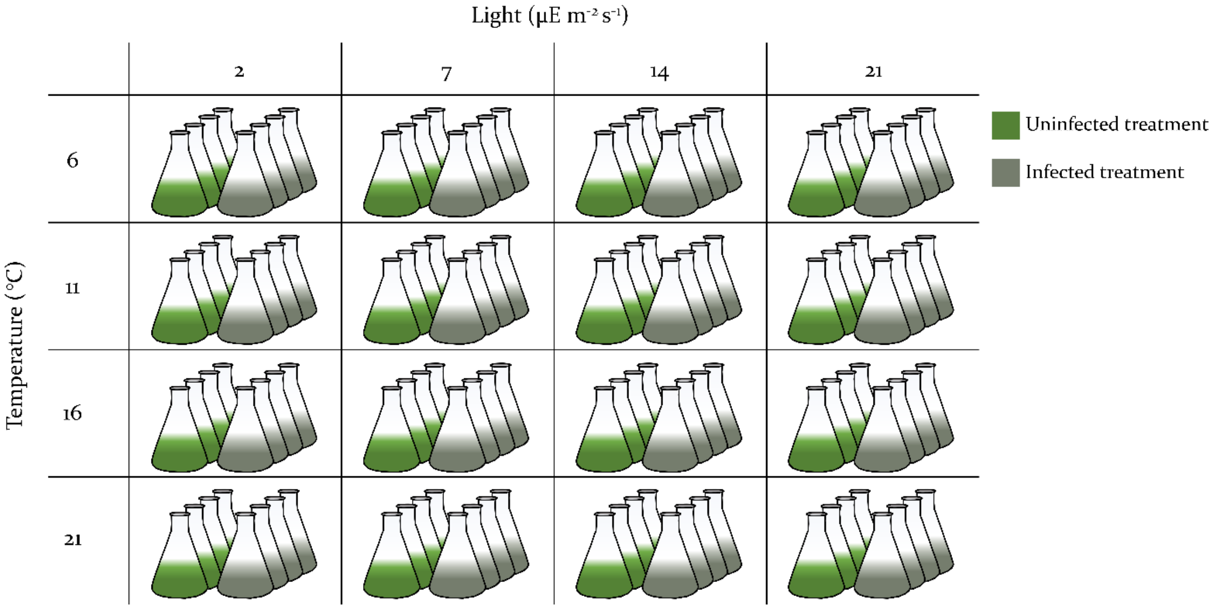


**Fig. S1**. Experimental setup indicating the factorial combination of temperature, light & infection levels, as well as the 4 replicates per treatment.

### *II. Model parameter dependence on temperature and light*

The infectivity parameter *I* is calculated as exponential decay of $H_u$ between $t_0$ and at *t* days (where t = 2). *Z* at $t_0$ is the initial zoospore concentration:

$$I = -\frac{ln\left[\frac{H_u(t)}{H_u(t_0)}\right]}{Z(t_0)\,t}$$

Fig. S2 illustrates how *I* changes with temperature and light, based on these calculations:

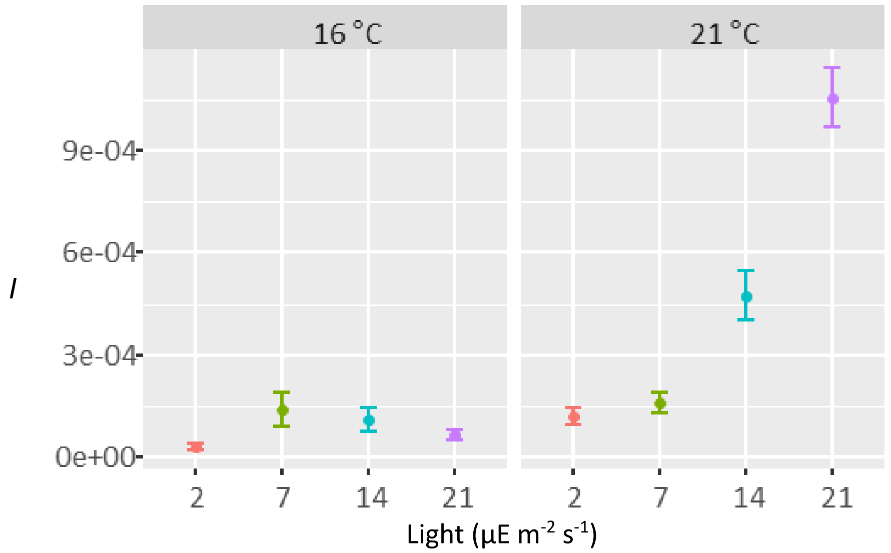


**Fig. S2**. Light and temperature-dependence of the infectivity parameter *I*

The production of zoospores *p* per nL bio-volume was calculated from supplementary data from Frenken et al. (2020). It is calculated from infected *P. rubescens* cultures (NIVA-CYA97/1). Production is the number of new zoospores divided by the bio-volume 2 days before. The zoospore production was calculated between days 4 to 10 when the infection is established, and prevalence of infection is still increasing.

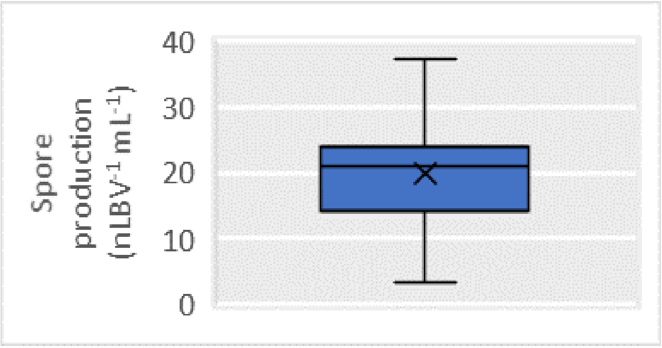


**Fig. S3**. Spore production per biovolume (nL $mL^{-1}$)

The model incorporates exponentially increasing or decreasing parameters with temperature for the development time of zoospores $\tau$, mortality rate of zoospores $m_z$, and the mortality rate of uninfected and infected host, $m_u$ and $m_i$. according to the following equation:

$$y = a \cdot e^{b \cdot T}$$

where *T* is temperature and *y* is the focal parameter.

Development time of zoospores $\tau$ has been set to 2 days at 21 degrees and 3 days at 16.
*a* is 16.42, *b* is -0.099.

Mortality rate of the uninfected host $m_u$ has been fitted to 1 percent of the respective growth rate.
*a* is 4.62e-04, *b* is 7.83e-02.

Mortality rate of the infected host $m_i$ is, like $m_u$, based on an exponential fit through the 1 percent growth rate, with an increase in the dependence on temperature of 50 percent, and a higher initial mortality rate.
*a* is 3.05e-02, *b* is 1.17e-01.

The life-time of zoospores was determined to be 2.71 days at 17 °C, which translates into a mortality rate $m_z$ of 0.36. subsequently the $m_z$ was set at 0.05, 0.12, and 0.5 for 6 °C, 11 °C and 21 °C respectively.
*a* is 0.035, *b* is 0.128.
